# The ALDH1A3 Reporter Construct: A Novel Mechanism of Identifying and Tracking the Breast Cancer Stem Cell Population

**DOI:** 10.1101/2023.09.14.557721

**Authors:** Nick Philbin, Ellen M. Laurie, Bre-Anne Fifield, Lisa Ann Porter

## Abstract

Cancer stem cells lie at the heart of progression and relapse for many solid tumours including breast cancers. The Breast Cancer Stem Cell (BCSC) population is typically isolated via a combination of markers utilizing various staining techniques which prevents the ability to track dynamic changes in expression and to dissect the role in pathogenesis overtime. Here we report the development of a reporter for the expression of Aldehyde Dehydrogenase 1A3 (ALDH1A3), a marker of high clinical importance in many breast cancers, and other solid tumours. BCSCs displaying increased transcriptional activation of ALDH1A3 demonstrate an increase in self-renewal capabilities. This tool improves the ability to reliably follow select cancer stem cell populations over time.

## INTRODUCTION

Breast cancer is a highly heterogenous disease with differences in genomic, proteomic, transcriptomic, and epigenomic landscapes contributing to not only inter-tumoural differences among patients but intra-tumoural differences leading to differences in characteristics inclusive of proliferation, stemness, migration, and invasion^1^. One source of heterogeneity is the high proportion of breast cancer stem cells (BCSC), a subset of the cancer cell mass contributing to intra-tumoural heterogeneity which retain features of mammary stem cells (Figure 1A)^2^. This is of clinical importance as BCSCs have the ability to not only self-renew but differentiate to recapitulate the heterogenous tumour mass^3,4^. Furthermore, these cells have increased chemoresistance and thus are thought of as the main drivers of relapse and secondary disease states^5,6^.

**Figure 1:**
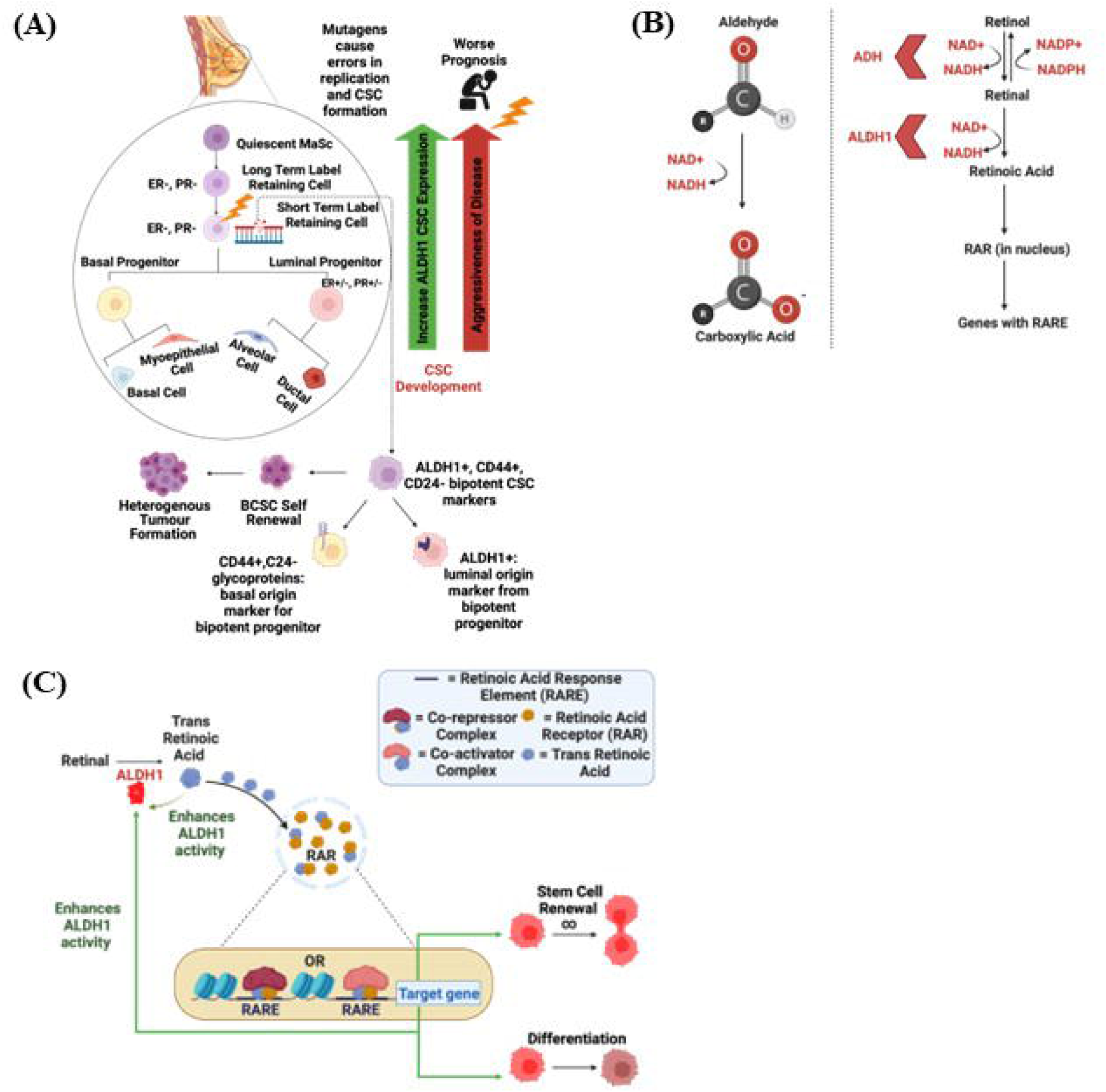
The Role of ALDH1 as a Prognostic Indicator and Its Implications on the Breast Cancer Stem Cell (BCSC) Population. **(A)** A visual representation of the breast stem cell hierarchy and the origin of breast cancer stem cells (BCSCs) with different potency. Mutations in cells with greater ALDH1+ expression contribute to the more aggressive cancers due to developing from a cell of higher origin. **(B)** A depiction of the oxidation of an aldehyde to a carboxylic acid; the generalized function of the cytosolic localized ALDH1. **(C)** The pathway associated with the production of retinoic acid and its downstream effects. Retinol is reversibly converted to retinal using alcohol dehydrogenase (ADH) and retinal is irreversibly converted to retinoic acid using aldehyde dehydrogenase (ALDH1). Retinoic acid has differential effects on the stem cell population depending on the gene it acts on leading to renewal or differentiation of cells within the tumour.

BCSC can be isolated and characterized by number of molecular markers. Typically, identification of these cell populations is achieved through cell surface marker expression via flow cytometry. Recently, Aldehyde Dehydrogenase 1 (ALDH1) has become the most clinically relevant markers for patient survival for select forms of breast cancer ^3,7,8^. ALDH1 is responsible for conversions of aldehydes to carboxylic acids through the oxidative process leading to subsequent downstream signaling contributing to stem cell renewal and differentiation (Figure 1B/C). There are 19 different isoforms of this enzyme. Interestingly, the ALDH1A3 isoform is proven to have the highest enzymatic activity and is amplified in 3.2% breast cancer cases with a greater proportion in TNBC^9^. This isoform is the main contributor of ALDH1 activity in breast cancer and is the most predictive of disease progression due to its involvement in a multitude of pathogenic processes including adhesion, migration, tumour growth, metastasis, and chemoresistance^10^.

Identification of the ALDH1 population is currently assessed using ALDEFLUOR™ assay. This assay measures endogenous ALDH1 enzymatic activity, detecting activity from 9 of the 19 ALDH1 isoforms (Figure 2A/B)^11^. A wealth of information has been obtained using this assay to dissect BCSC dynamics and highlights the importance of ALDH1 expression in the BCSC population. A limitation of this assay is that it does not allow for monitoring of this population in real time, or isolation of the role of specific ALDH1 isoforms. More specific methods of detecting ALDH1 isoforms are needed to better monitor ALDH1 population dynamics and the implications on tumour evolution and patient prognosis.

**Figure 2:**
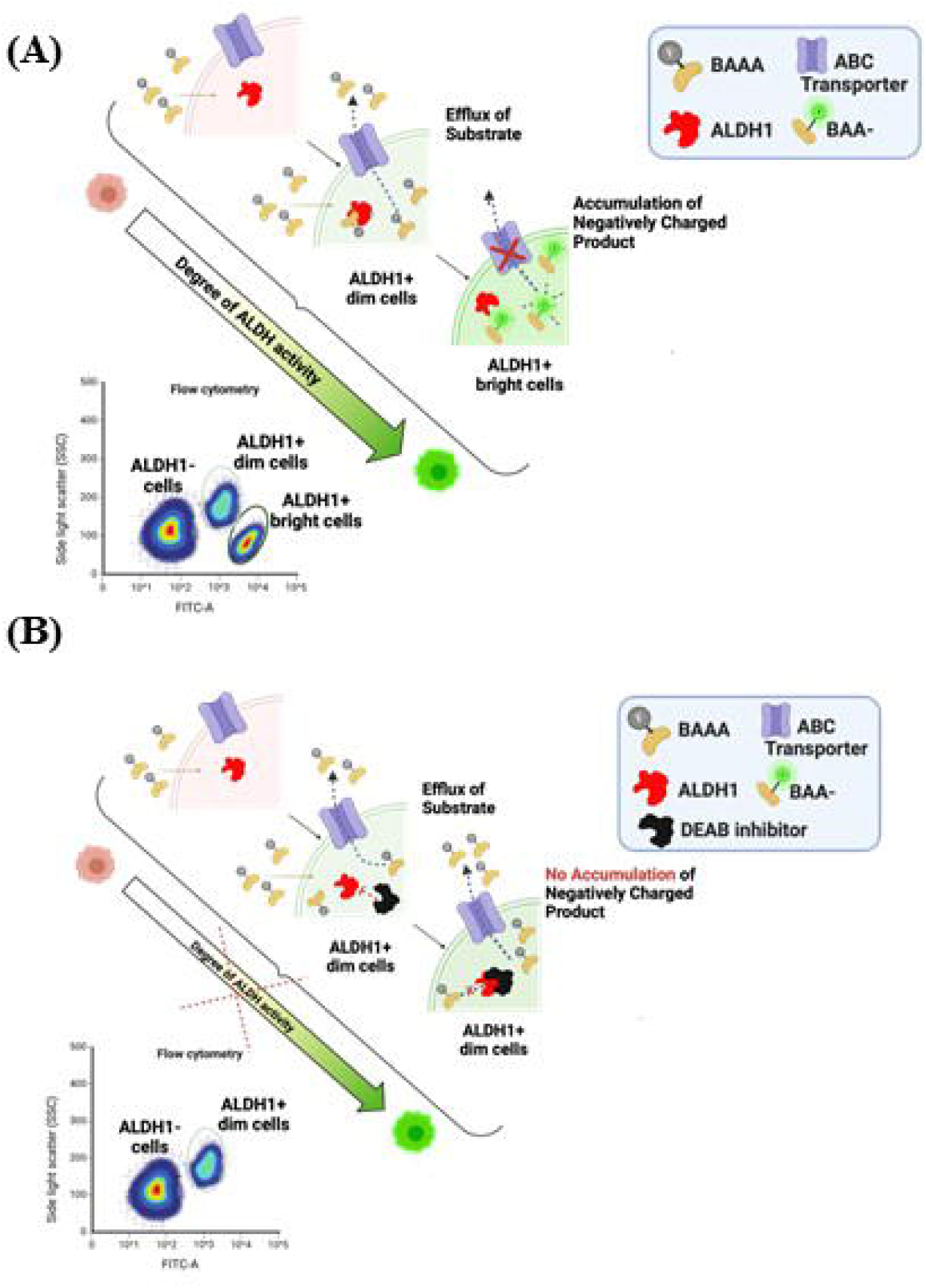
The Mechanistic Basis behind the ALDEFLUOR™ Assay’s ability to detect ALDH1 activity. **(A)** ALDH1 acts to break down the fluorescent aldehyde BODIPY-aminoacetylaldehyde (BAAA) to BODIPY-amino acetate (BAA-). The negatively charged product accumulates in the cytoplasm and its fluorescent emission allows these cells to be detected by flow cytometry with the greater the accumulated product, the greater the fluorescence emission (**B)** A selective inhibitor for ALDH1 known as N,N-diethylamino benzaldehyde (DEAB) is used as a negative control due to its ability to prevent the conversion to the fluorescent BAA- and subsequent accumulation in the cytosol.

We developed an ALDH1A3 reporter construct where the promoter and enhancer regions of ALDH1A3 drive expression of a tdTomato fluorophore. Importantly, like the ALDH1A1 gene of which reporters have already been developed, it is regulated transcriptionally and thus serves as an accurate method of detection^12–14^. We show that this reporter construct is able to accurately isolate ALDH1A3 positive BCSC populations and this correlates with fluorescent signal intensity. This tool lends itself the opportunity to be utilized *in vivo* to track ALDH1A3+ population dynamics overtime and its role in processes contributing to tumour progression.

## MATERIALS AND METHODS

### Cell Culture

MDA-MB-231 (Triple Negative Breast Cancer (TNBC)) and MCF-7 (Estrogen Receptor Positive (ER+)) cell lines (ATCC-HTB-26 and HTB-22) were maintained in full growth media consisting of Dulbecco’s Modified Eagle’s Medium (DMEM) High Glucose, 10% Fetal Bovine Serum (FBS;Gibco), and 1% Penicillin-Streptomycin (P/S; Invitrogen). Cells were cultured once they reached 70-80% confluency. Cells were subcultured using 0.25% trypsin. Cells were counted using a trypan blue exclusion assay. All cell lines were maintained at 37°C and in a 5% CO_2_ environment.

### Mammosphere Formation Assay

Cells were seeded at a density of 10,000 cells /mL of DMEM-F-12 (Corning 10-092) supplemented with 20ng/mL epidermal growth factor (Gibco; PHG0315), 20ng/mL human fibroblast growth factor (Sigma F5392), B27 (Thermofisher Scientific; 17504044), 0.1mg/mL gentamicin (Gibco; 15710-064), 5μg/mL heparin (Sigma; H0777) and 1% P/S in ultra-low attachment plates. Mammospheres were dissociated with 0.05% trypsin to obtain single cell suspensions and seeded at a density of 2500 cells/mL for secondary sphere formation. Quantification of mammosphere diameter was performed via ImageJ software. The total number of spheres ≥ 40μm were quantified and divided by the number of cells seeded and multiplied by 100 to gain a percentage: mammosphere formation efficiency (% MFE).

### Viral infection

10,000 cells were seeded in a 96 well plate. After 24 hours, cells were infected in serum and antibiotic free media containing 8μg/mL of polybrene (Santa-Cruz; SC134220) and lentivirus at an MOI of 3. Cells were incubated for 24 hours and viral media was removed and replaced with full growth media.

### ALDEFLUOR™Assay

ALDH1 isoforms were detected using the ALDEFLUOR™ assay (STEMCELL Technologies; 01700) as per manufacturer’s instructions. MDA-MB-231 and MCF7 cells were collected and suspended in a volume representative of one million cells/mL of ALDEFLUOR™ assay buffer. Cells were administered 5μl of activating reagent per million cells and subsequently half of the reaction volume was added to a tube with the ALDH inhibitor, N,N-diethylaminobenzaldehyde (DEAB). Tubes were incubated for between 20-45 minutes at room temperature (RT) or 37°C. Cells were subsequently spun down, resuspended in 500ul of ALDEFLUOR™ assay buffer, stored on ice and ran on the flow cytometer. Each inhibited sample (tube with DEAB constituent) was analyzed and gated prior to the uninhibited sample.

### Plasmid Construction

pHIV-tdTomato (Addgene plasmid #21374) was constructed by Dr. Bryan Welm. This plasmid was utilized to construct the pHIV-ALDH-1A3-tdTomato reporter construct used for lentiviral infection. Amplification of the ALDH1A3 isoform genomic promoter and enhancer region (NCBI Blast: **NG_012254.1**, chromosome 15, bp 4173-5104) by PCR reaction (primers shown in Table 1) was performed. This region along with the pHIV-tdTomato plasmid was cut with the restriction enzymes AgeI (BshTI) and XbaI. The endonuclease cut vector and the ALDH1A3 promoter and enhancer region were run on a 1% agarose A gel and extracted via the Monarch Gel Extraction Kit (Cat#T1020S) from New England BioLabs. The ALDH1A3 promoter and enhancer region were then ligated to the vector backbone using 5units/μl T4 DNA ligase to replace the elongation factor-1 (EF-1) promoter (Figure 3).

**Table 1:**
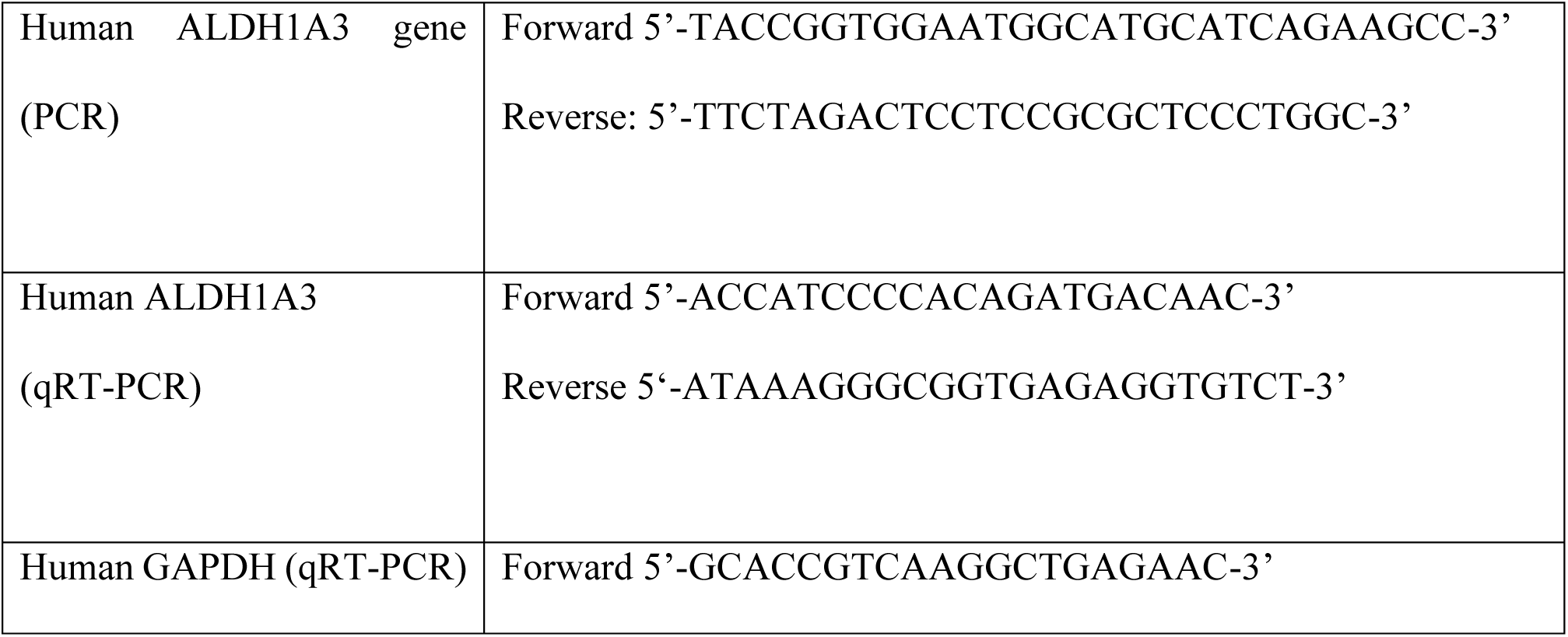
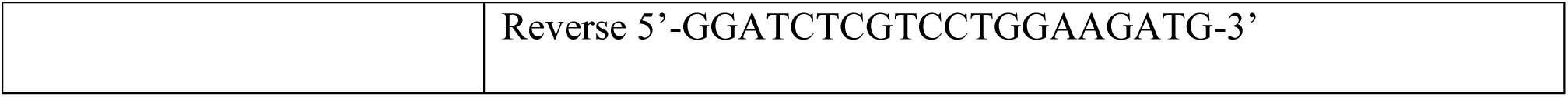
Primers Utilized for PCR and qRT-PCR.

**Figure 3:**
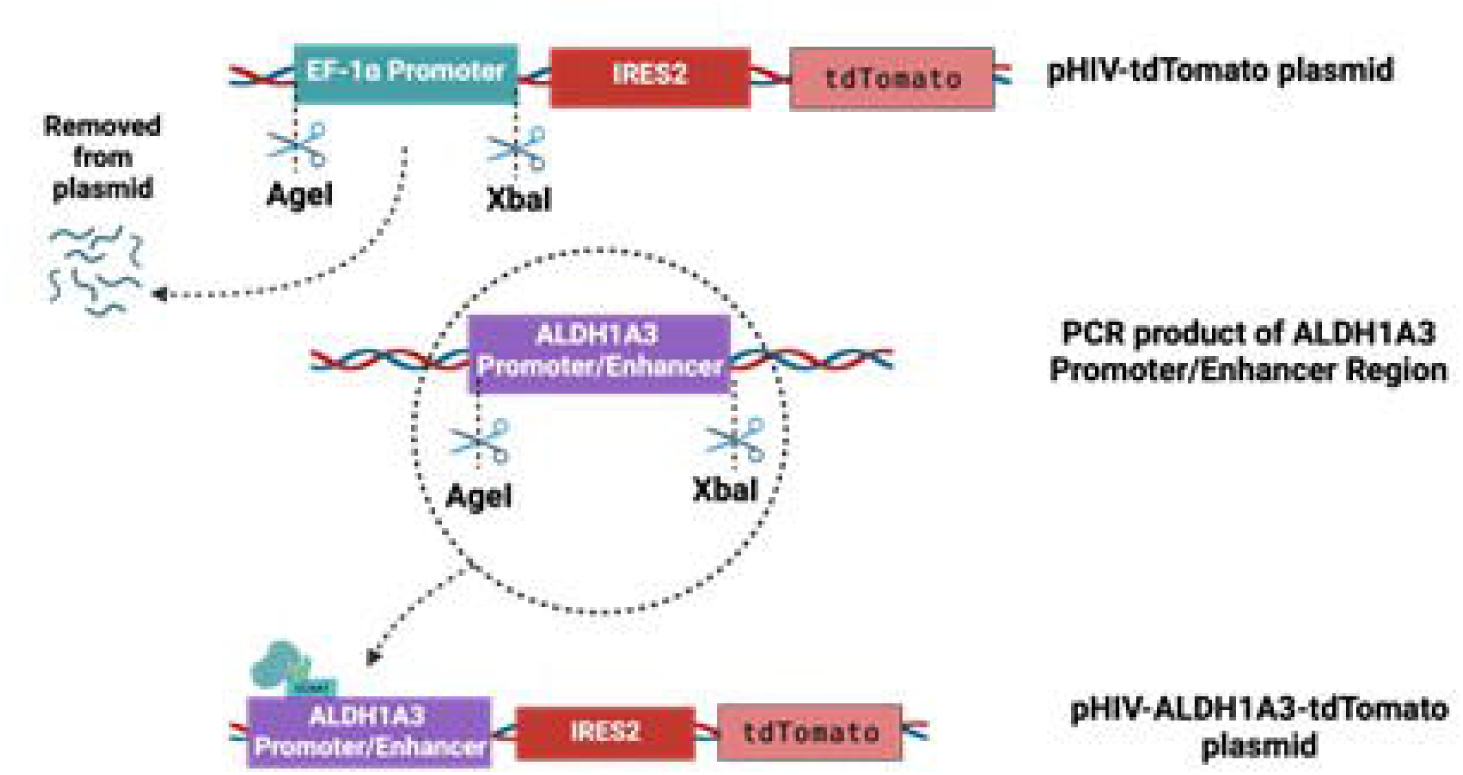
The Mechanism Behind the Creation of the pHIV-ALDH-1A3-tdTomato reporter system. The pHIV-tdTomato plasmid and ALDH1A3 promoter and enhancer regions were cut with AgeI (BshTi) and XbaI. Digest of the pHIV-tdTomato plasmid led to the removal of the EF-1 promoter. Subsequently, the ALDH1A3 promoter and enhancer region was ligated into the pHIV-tdtomato backbone leading to the construction of the aforementioned ALDH-1A3-tdTomato construct.

### Flow Cytometry

Cells were collected and resuspended in 500μl of BD Pharmigen Stain Buffer (BD Biosciences; 554656), the ALDEFLUOR™ Buffer or Hank’s Balanced Salt Solution (Thermofisher Scientific; 24020-117) with 2% FBS (HBSS+2%FBS) before being run the BD LSR Fortessa X—20 at the University of Windsor Flow Cytometry Facility.

### Fluorescent Activated Cell Sorting (FACs)

Cell lines were collected and resuspended in resuspended in a solution of 1x Phosphate Buffer Saline (PBS) + 1% FBS+ 2mM Ethylenediaminetetraacetic acid (EDTA) at a density of 1 million cells/mL. Cells were then filtered through a 70μm strainer and run on the BD FACSARIA Fusion™ at the University of Windsor Flow Cytometry Facility.

#### ALDEFLUOR™ Assay Gating

Cells were gated into one of two populations: ALDH1- and ALDH1+. Cells negative for ALDH1-were deemed to be less than 10^3^ on the logarithmic scale for FITC. For each experiment, the unstained control was gated first, followed by the DEAB inhibited sample, and then the experimental sample. The percentage of cells deemed to be ALDH1+ was calculated as the difference between the ALDH1+ positive cells for the experimental sample minus the ALDH1+ cells for the inhibitor sample.

#### ALDH1A3 Reporter Gating

Cells were gated into one of four populations: ALDH1A3 negative (ALDH1A3-), ALDH1A3+ low, ALDH1A3+ medium, or ALDH1A3+ high on a logarithmic scale for PE-Cy5. The ALDH1A3-population was defined as a fold change of less than 10^3^, ALDH1A3+ low population as between 10^3^ and 10^4^, ALDH1A3 medium as halfway between 10^4^ and 10^5^, and high as above the halfway point between 10^4^ until 10^5^. Percentage of ALDH1A3+ positive cells was calculated from the number of positive cells in each gate divided by the total number of live cells in the sample, multiplied by 100.

### Quantitative Real-Time Polymerase Chain Reaction (qRT-PCR)

RNA extractions were performed using the Qiagen RNeasy Plus Mini Kit (Qiagen; 74104). cDNA was synthesized using Quanta Q Script master mix (VWR; CA101414-104). qRT-PCR was performed using SybrGreen detection via the Fast SYBR™ Green Master Mix (Thermofisher; 4385612). 10μl reactions were run for forty cycles on the Viia7 Real-Time PCR System and Software (Life Technologies). Samples were normalized to GAPDH and analysis performed via the QuantaStudio™ Real-Time PCR Software. Primers utilized were as shown in Table 1.

## RESULTS

### ALDH1A3+ Populations show Increase in Transcript levels Corresponding with Increase in Fluorescent Intensity

The ALDH1A3 gene promoter and enhancer region were amplified and cloned into the pHIV-tdTomato. Breast cancer cell lines known to have differing expression of ALDH1A3 were used for validation. The MCF7 ALDH1A3+ population encompasses less than 1% of cells compared to approximately 10% in the MDA-MB-231 cell line, however results have varied between different groups^3,15,16^. MDA-MB-231 and MCF7 cells were infected with ALDH-1A3-tdTomato construct, and sorted into ALDH1A3-, ALDH1A3+ low, ALDH1A3+ med, and ALDH1A3+ high populations. Cells were subsequently collected and subject to qRT-PCR analysis for ALDH1A3 transcript levels (Figure 4A). Relative to the ALDH1-population; MDA-MB-231 transcript levels increased 133%, 161%, and 199% for the ALDH1A3+ low, medium and high populations (Figure 4B) and 334%, 596%, and 937% for the MCF7 cell line (Figure 4C). Intensity of fluorescence correlates with ALDH1A3 transcript expression, confirming the gating strategy and validity of this reporter in isolating distinct populations based on ALDH1A3 expression.

**Figure 4:**
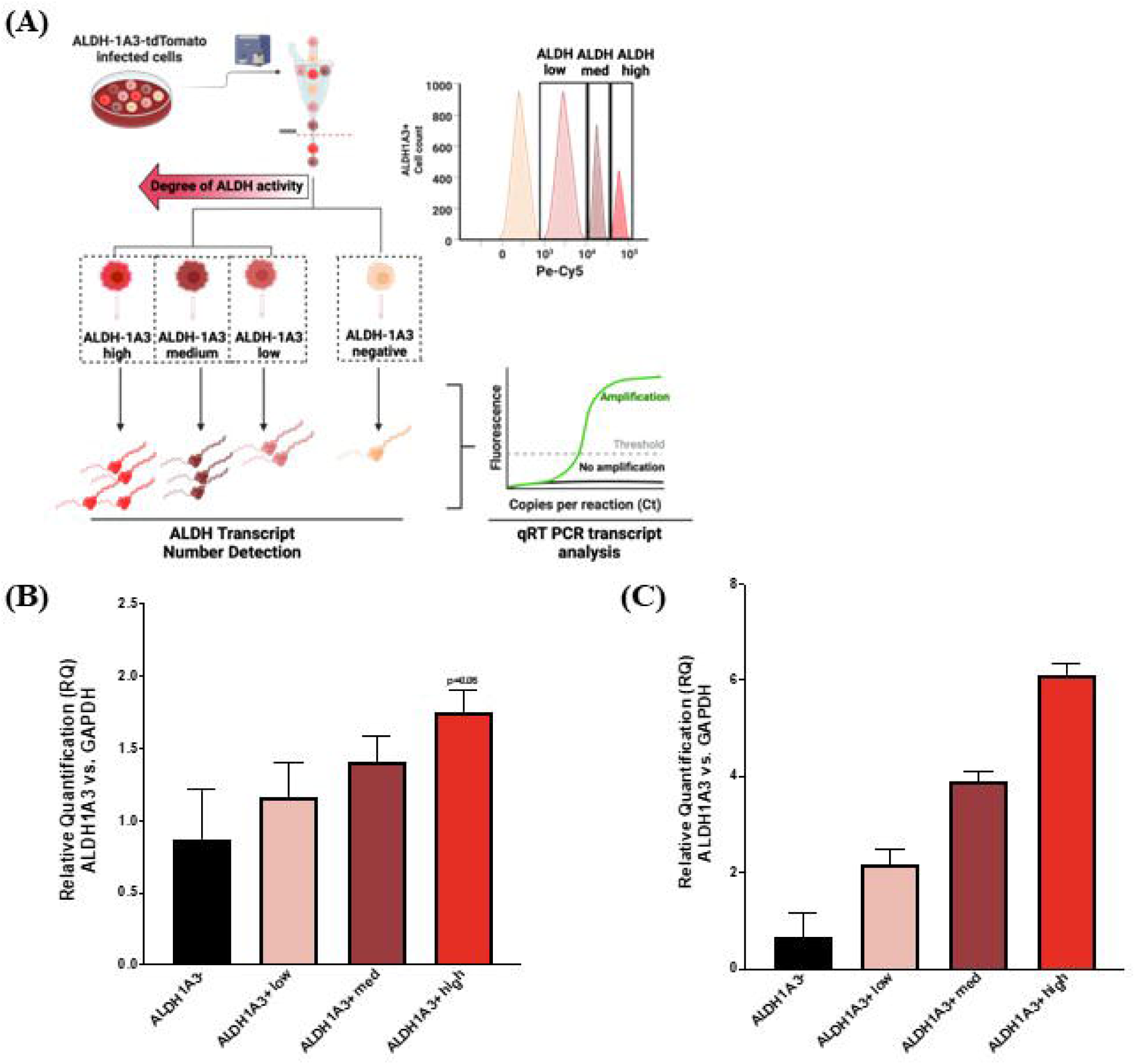
Validation of the ALDH1A3 reporter system through transcript analysis. **(A)** MDA-MB-231 and MCF7 cells were infected with the pHIV-ALDH-1A3-tdtomato reporter construct, run on the flow cytometer and sorted into ALDH-1A3 low, medium and high populations. ALDH-1A3 transcript levels of the low, medium, and high gated populations via qRT-PCR analysis for **(B)** MDA-MB-231 cell line and **(C)** MCF7 cell line relative to GAPDH control. Students t-test, Error bars represent standard error (SE); n=3.

### ALDH1A3 Increase in Transcript Levels Correlate with an Increase in Sphere Formation Efficiency

Primary and secondary sphere formation assays were conducted using the gated populations described above to assess the self-renewal capabilities of each of the 4 populations and determine if it correlated with ALDH1A3 transcript levels (Figure 5A). Primary sphere formation efficiency for the MDA-MB-231s was 0.2%, 1.32%, 1.70%, and 2.93% for the ALDH1-, ALDH1A3+ low, ALDH1A3+ medium, and ALDH1A3+ high populations respectively (Figure 5B) and 0.10%, 0.33%, 0.57%, and 1.09% for the MCF7 cell populations. (Figure 5C). Secondary sphere formation efficiency was 0.57%, 4.05%, 5.34%, and 11.75% and 0.21%, 0.35%, 0.72%, and 1.25% for the MDA-MB-231 (Figure 5B) and MCF7 cells respectively for the previously mentioned populations (Figure 5C). Thus, increasing sphere forming efficiency correlates with increasing ALDH1A3 transcript expression.

**Figure 5:**
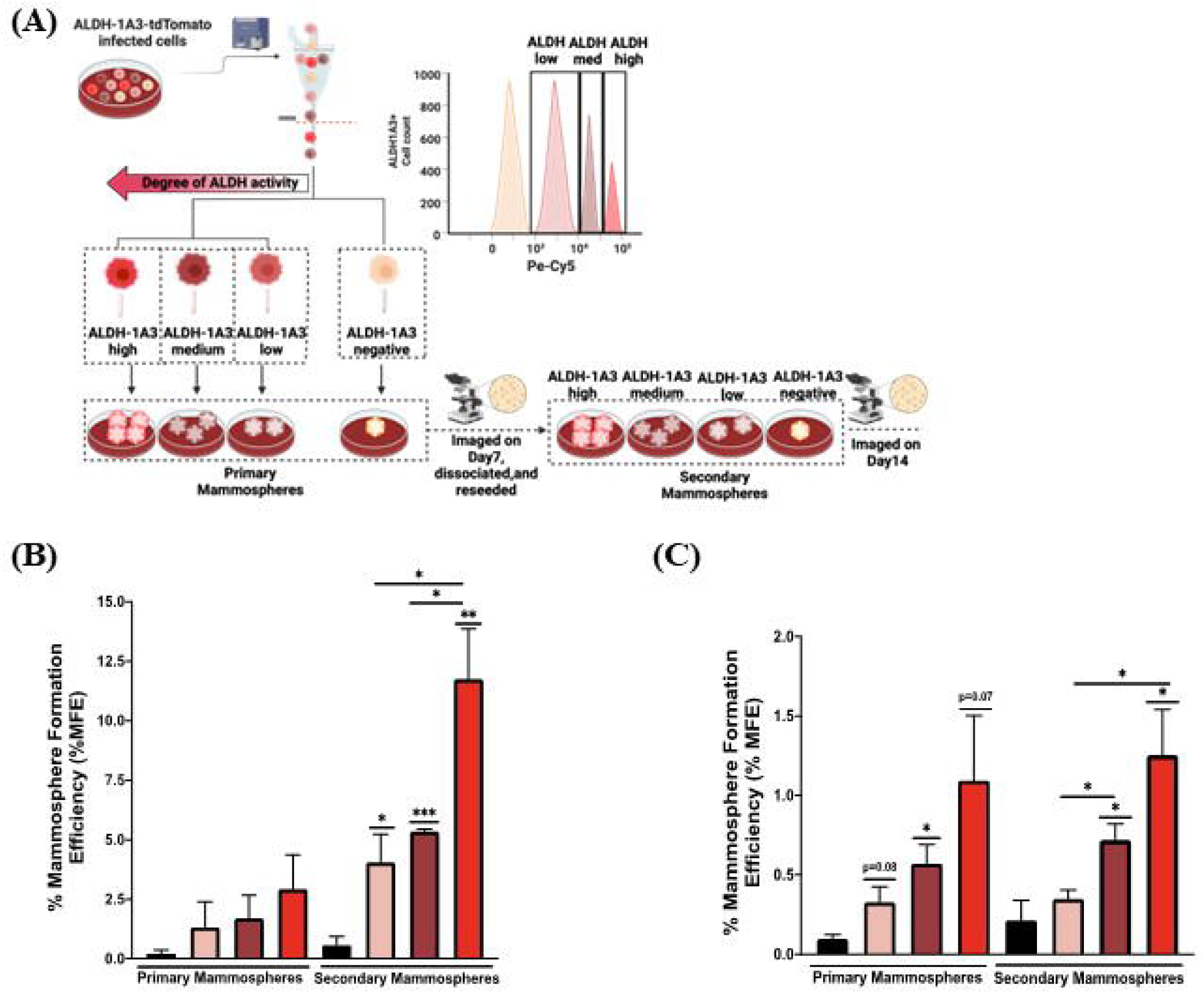
Validation of the ALDH1A3 reporter system through a functional output assay to determine stem-like potential. **(A)** MDA-MB-231 and MCF7 cells were infected with the pHIV-ALDH-1A3-tdtomato reporter construct, run on the flow cytometer and sorted into ALDH-1A3 low, medium and high populations. Cells from each population were then seeded for a mammosphere formation assay. Results were represented as the %MFE of ALDH1A3+ low, medium, and high populations for **(B)** MDA-MB-231 cell line and **(C)** MCF7 cell line. Quantification of spheres >40μm in diameter. Students t-test, p*<0.05, p**<0.01, Error bars represent SE; n=3.

### ALDH-1A3 Reporter Demonstrates Minimal Variability in ALDH1A3 Population Detection in a Single Heterogenous Population

The pHIV-ALDH-1A3-tdTomato reporter construct was infected into MDA-MB-231 and MCF7 cells, grown up, seeded at a cell density of 300,000 cells on a 10cm plate, allowed to grow for two days and subsequently split into three tubes and run on the flow cytometer to assess the consistency of ALDH1A3 detection within a given heterogeneous population (Supplementary Figure 1A/B). The three samples had an average ALDH1A3+ population of 31.7%, 46.9%, and 10.1% for the low, medium and high populations respectively. Samples proved to vary on average within a given infection by 0.15%, 0.31%, and 0.06% for the ALDH1A3+low, medium, and high populations respectively across three samples for the MDA-MB-231s (Supplementary Figure 1C). For the MCF7s, the average ALDH1A3+ population was 33.8%, 41.9%, and 14.2% for the low, medium and high populations respectively. Samples only varied by 0.27%, 0.51%, and 0.23% for said populations (Supplementary Figure 1D). An analogous experiment was performed utilizing the ALDEFLUOR™ assay where a sample was administered the activating agent and half was utilized for the inhibitor control tube. Subsequently, the remaining portion of the reaction was split into three and ALDH1+ activity was measured via flow cytometry (Supplementary Figure 1E). The MDA-MB-231 cell line had an average cell positivity of 0.001% and a variability of 0.009% within a sample (Supplementary Figure 1F). Meanwhile, the MCF7 cell line had an average cell positivity of 0.18% and a variability of 0.06% within samples (Supplementary Figure 1G). In each case, an increase in variability was seen compared to the aforementioned reporter.

### Breast Cancer Cell Lines Show Distinct Patterns of ALDH1A3+ Cell Population Dynamics with High Replicability Between Infections

Following validation of the pHIV-ALDH-1A3-tdTomato reporter to accurately measure ALDH1A3+ activity in the MDA-MB-231 and MCF7 cell lines, it was used to characterize the ALDH1A3+ activity in two additional cell lines; the MDA-MB-453 and the MDA-MB-468s, across three separate infections (Figure 6). When combining this data with the original cell lines used to validate the reporter, the MDA-MB-231 and MDA-MB-453 cell lines had the highest percentage of ALDH1A3+cells in the low activity population, followed by medium, and then high. While the MCF7 cells has the greatest percentage of ALDH1A3+cells in the medium activity range, followed by low and then high. The MDA-MB-468 cell line only showed a relatively low percentage of positive cells, with all being in the low population. Across three infections for each cell line, the variability was low for the ALDH1A3+ low, medium and high cell populations (respectively) for the cell lines which follow. The MDA-MB-231 cell line had a variability of 1.39%, 2.83%, 1.05%. The MCF7 a variability of 3.09%, 2.18%, 2.06%. The MDA-MB-453 a variability of 1.99%, 1.32%, 0.033% and the MDA-MB-468 a variability of 1.22%. Notably, this was considerably more consistent than the commercially available ALDEFLUOR™ KIT where samples varied a minimum of ∼38% between repeats and even more dramatically between different protocols described in the literature (Supplementary Figure 2).

**Figure 6:**
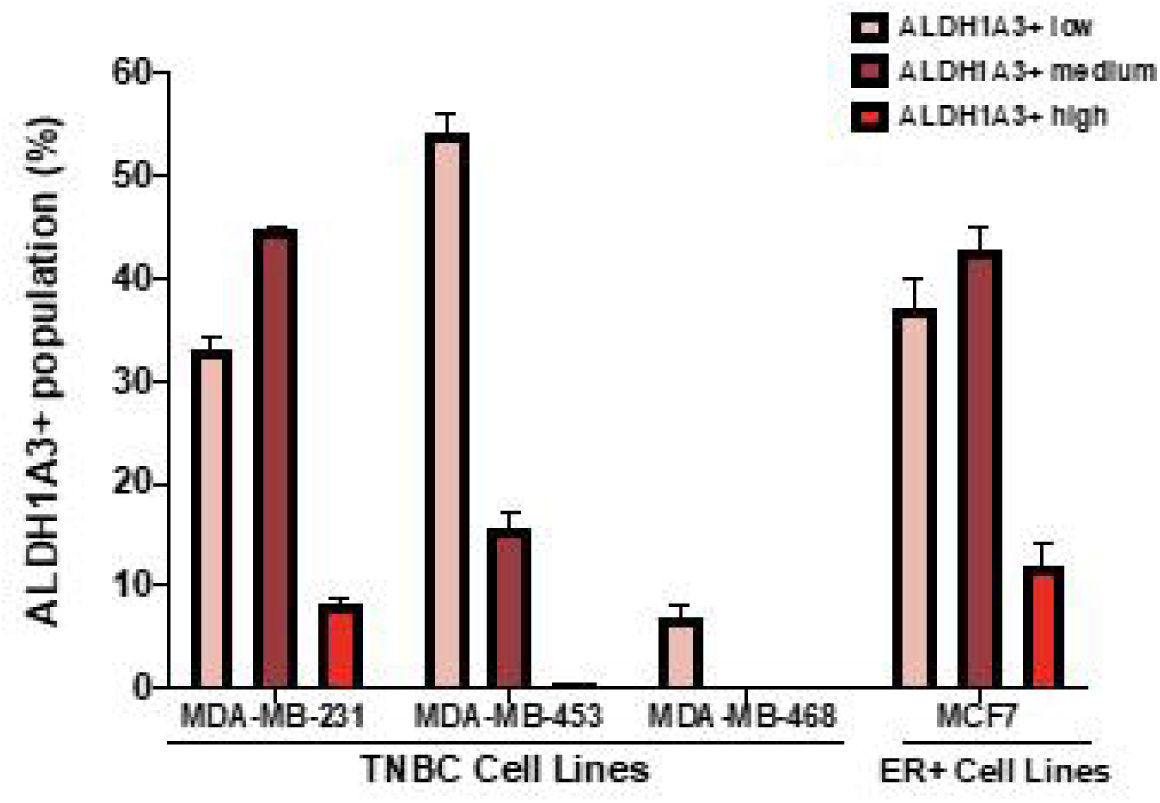
Characterization of Endogenous ALDH1A3+ activity in a panel of Breast Cancer Cell Lines. The quantification of the percentage of cells determined to be ALDH1A3+ via flow cytometry. ALDH1A3+ cells are represented in terms of the percentage of the total cell population. Error bars are representative of the standard error of the mean. Student’s t-test, Error bars represent SE; n =3.

## DISCUSSION

The ALDH1A3 reporter construct described in this work allows for a specific and reliable method of tracking cancer stem cells. Our data shows that cells which have higher fluorescent emission, correspond to an increase in ALDH1A3 transcript production, which is indicative of stemness based on its regulatory mechanisms lying at the expression level^17–24^. This increase in expression led to an increase in sphere formation efficiency, which measures the self-renewal capabilities of a cell population. This aligns with the fact that ALDH1A3 overexpression has been shown to impart increased colony formation in MDA-MB-231 and MDA-MB-468 cell lines used for these assays ^25^. It is important to note that, across the four cell lines that were characterized, the percentage of ALDH1A3+ high cells mimicked what is reported in the literature with the MDA-MB-231s having a low ALDH1A3+ high comparable to the MCF7 and greater than the MDA-MB-453^26,27^. Of note, the MDA-MB-468s had the lowest percentage of positive cells compared to the other cell lines. Importantly, conflicting results in terms of the ALDH1+ population percentage have been reported via the ALDEFLUOR™ assay reporting low ALDH1 activity as around 10% of cells and others as above 80% for this cell line^15,26,28,29^. Additionally, the MDA-MB-468 cell line has been shown to have reduced ALDH1 activity compared to the MDA-MB-231 and MCF7 cell lines via immunofluorescence assays^30^. Reducing variability is an important concern and our results support that this reporter provides more reliability between assays.

This reporter measures a single isoform (ALDH1A3) as compared to the potential nine isoforms which could contribute to the phenotype seen if using the ALDEFLUOR™ assay leading to a more specific interpretation of data and reducing the variability seen with the ALDEFLUOR™ assay within a given cell population. This is extremely important for not only breast cancer but other cancers such as ovary, skin, liver, brain, pancreatic and esophageal which also heavily rely on the 1A3 isoform expression^31–36^.

An additional benefit of this system is the experimental ease of which it can be used and the lack of optimization needed to detect ALDH1 activity. The ALDEFLUOR™ assay has proven the need to optimize both time and temperature conditions in a cell line specific manner and the ALDH1 population dynamics have yet to be characterized in many cases.

A third benefit of this system is the ability to clearly distinguish the magnitude of ALDH1 activity. With our reporter, the degree of fluorescence is directly correlated to the binding of the activating complex to the promoter/enhancer regions triggering the initial steps of transcription. Meanwhile, in the ALDEFLUOR™ assay, the substrate which is added to the reaction mixture, is already dimly fluorescing which can contribute to additive effects to overall activity being detected if not fully inhibited, as it has been noted that the DEAB inhibitor is not capable of inhibiting most ALDH1 isoforms completely^31^. All of these are important for stem cell number identification due to their extreme potency. The reporter construct depicts a more widespread gradient of ALDH1A3 activity detected compared to the ALDEFLUOR™ assay. This allowed for the ability to detect a cell population with medium expression of ALDH1A3 in addition to the low and high populations which are isolated with the ALDEFLUOR™ assay. This could give insight into a third cell population aside from the ALDH1 low and bright isolated populations via the ALDEFLUOR™ assay and their role in BCSC mediated processes.

The last benefit of our reporter system is the replicability and minimal variability in marking the ALDH1A3 population both between and within infections. This could be due to the fact that the ALDEFLUOR™ assay is picking up additional isoforms in addition to the 1A3 isoform leading to a greater ALDH1+ population difference between different cell populations. This may be in part why the inhibitor control tube can result in a greater ALDH1+ cell population detected in comparison to the experimental sample from a single heterogenous population as seen in certain optimization experiments. In addition, the DEAB inhibitor has been demonstrated to serve as a substrate for certain ALDH1 isoforms, albeit a slow one for isoforms such as ALDH1A3, but when dealing with such a small population of positive cells, it may be a factor^37^.

It is important to note that one marker is not enough to denote the potency and tumourigenic effects of a given cancer and combinatorial analysis provides a more robust and greater clinical indicator of patient prognosis. Cells determined to be CD44+/CD24-/ALDH1+ have been most indicative of tumourigenesis and poor patient outcomes^38^. A study done in NOD-SCID mice demonstrated that as few as 20 CD44+/CD24-/ALDH1+ cells were able to potentiate tumour growth upon mammary fat pad transplantation as compared to 1500 ALDH1+ cells, and upwards of 50,000 CD44-/CD24-/ALDH1-cells which were unable to form tumours at all^3^. In addition, tumours with high CD44+/CD24-/ALDH1+ have been demonstrated to have increased incidences of distant metastasis^38^.

That being said, reporter constructs for proteins such as CD44 are difficult approaches for tracking expression levels in single cells due to regulation occurring post-transcriptionally and the output function being dependent on spliced variant production, and ligand specificity^39^. Our reporter construct could be used in combination with staining approaches following ALDH1A3+ cell isolation or the previously established pluripotency stem cell marker reporter constructs for proteins such as OCT4, SOX2, NANOG and ABCG ^40–42^. This combination approach may improve the detection of CSC populations and the determination of their role during tumour progression and evolution ^41^. Together, this data describes a reporter system capable of tracking CSCs expressing ALDH1A3. We position this as a tool to be used for providing insight into the biology of pathogenesis by the CSC population and the implications on phenomenon such as the increased frequency of relapse in this population.

## LIST OF ABBREVIATIONS

ADH: Alcohol Dehydrogenase
ALDH1: Aldehyde Dehydrogenase One
BAAA: BODIPY-aminoacetylaldehyde
BAA-: BODIPY-amino acetate
BCSC: Breast Cancer Stem Cells
CSC: Cancer Stem Cells
DEAB: N,N-diethylaminobenzaldehyde
EDTA: Ethylenediaminetetraacetic acid
EF-1: Elongation Factor One
FBS: Fetal Bovine Serum
GAPDH: Glyceraldehyde 3-phosphate dehydrogenase
HBSS: Hanks Balanced Salt Solution
MFE: Mammosphere Formation Efficiency
PBS: Phosphate Buffer Saline
TNBC: Triple Negative Breast Cancer

## AUTHOR CONTRIBUTIONS

N-P, E-ML, and B-F carried out the execution of the experiments. LAP funded the project and had a lead role in study design, interpretation of the data and manuscript preparation. N-P, B-F and LAP prepared the manuscript and provided comments on the intellectual content. All authors approved the final manuscript.

## ACKNOWLEDGEMENTS

We would like to thank Dr. Bryan Welm for the gift of the pHIV-tdtomato plasmid (Addgene plasmid #21374) which was used to clone the ALDH1A3 reporter constructs. We would like to thank Dr. Elizabeth Fidalgo Da Silva, manager of the UWindsor Flow Cytometry Facility for her assistance.

