## Supplemental Figures for "The ALDH1A3 Reporter Construct: A Novel Mechanism of Identifying and Tracking the Breast Cancer Stem Cell Population"

**Supplementary Figure 1: The ALDH1A3 Reporter Construct Infections are Replicable and Demonstrate Greater Consistency in Characterization Within a Heterogenous Cell Population than the ALDEFLUOR$^{TM}$ Assay. (A)** Cells were infected with the pHIV-ALDH-1A3-tdTomato construct grown up, seeded at a density of 300,000 cells on a 10cm plate and allowed to grow for two days. Cells were then split into 3 tubes per infection before being run on the flow cytometer. **(B)** Data is gated and shown as % ALDH1A3+ low, medium, and high of the entire cell population represented as P2, P3, and P4 respectively for the **(C)** MDA-MB-231 and **(D)** MCF7 cell lines. **(E)** 300,000 cells were seeded on a 10cm plate, collected two days later and stained with the ALDEFLUOR$^{TM}$ Kit before being split into three for being run on the flow cytometer. Quantification of the ALDH1+ cells relative to the entire cell population for the **(F)** MDA-MB-231 and the **(G)** MCF7 cell lines respectively. Error bars represent SE; n=3.

**Supplementary Figure 2: Optimization of the ALDEFLUOR$^{TM}$ Kit for Endogenous ALDH1 Detection Yields Differences in Population Dynamics. (A)** A single plate was collected and split in two tubes, which were subsequently split into two further tubes. The tubes were then administered the activating agent alone or in combination with the DEAB inhibitor and run on the flow cytometer. **(B)** Representative flow plots used to gate the ALDH1+ population where P2 and P3 represent the ALDH1- and ALDH1+ populations respectively. **(C)** Quantification of ALDH1+ cells identified from the protocol in panel A. Samples varied in ALDH1+ cell identification by 38.5% within a sample and 38.9% between samples. **(D)** MDA-MB-231 and MCF7 cell lines were seeded on 10cm plates at a 300,000 cell density and collected two days later and subsequently stained with the **ALDEFLUOR$^{TM}$ Kit** for 20 minutes at 37°C, 40 minutes at room temperature or 40 minutes at 37°C before being run on the flow cytometer. Quantification of the ALDH1+ cell population for the **(E)** MDA-MB-231 and **(F)** MCF7 cell lines relative to the entire cell population. Student's t-test, Error bars represent SE; $p^* < 0.05$, $p^{**} < 0.01$, $p^{***} < 0.001$, (n=2, n=3).

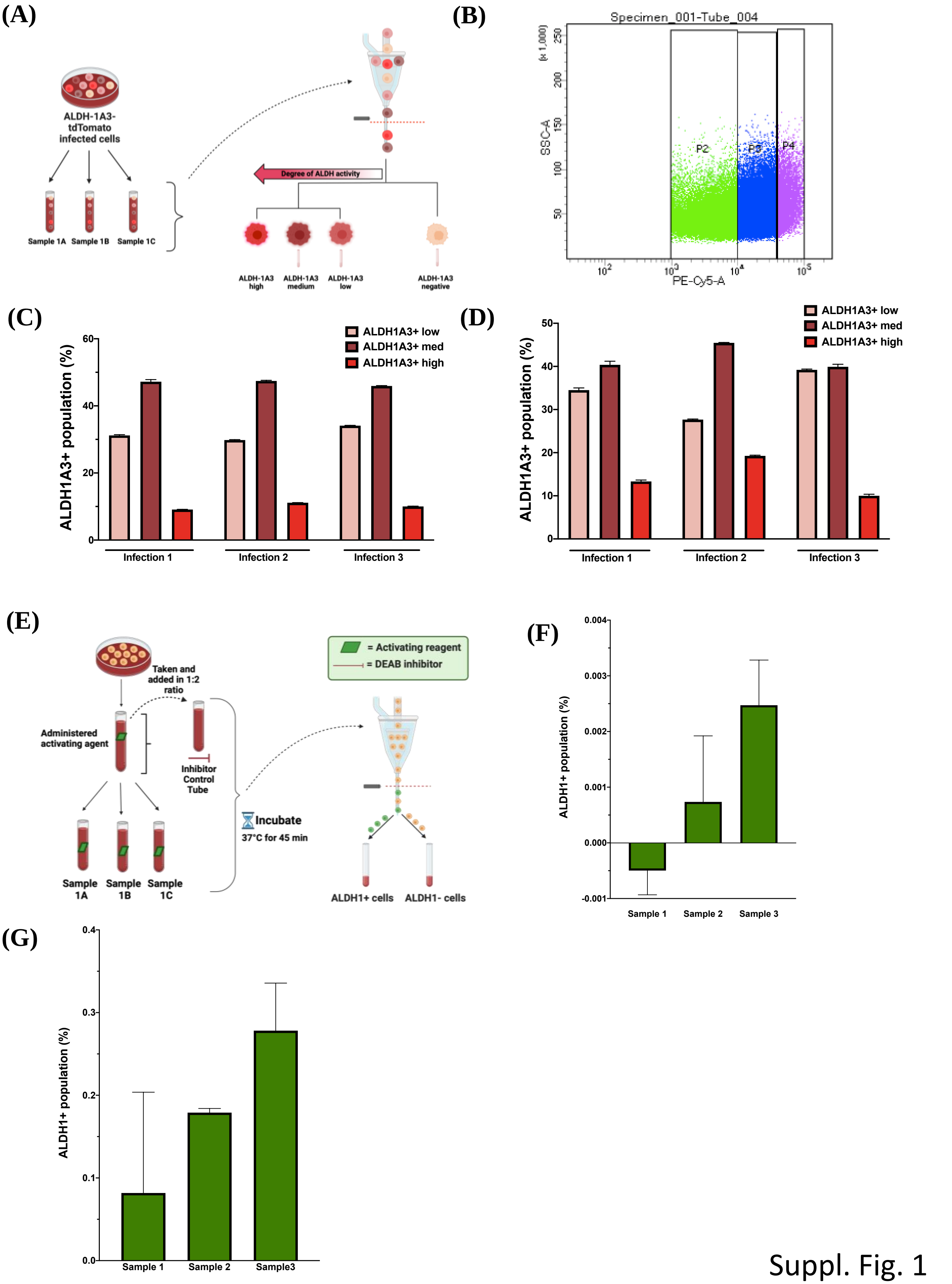
(A)
ALDH-1A3-tdTomato infected cells
Sample 1A
Sample 1B
Sample 1C
Degree of ALDH activity
ALDH-1A3 high
ALDH-1A3 medium
ALDH-1A3 low
ALDH-1A3 negative
(B)
Specimen_001-Tube_004
SSC-A (x 1,000)
P2
P3
P4
PE-Cy5-A
(C)
ALDH1A3+ population (%)
ALDH1A3+ low
ALDH1A3+ med
ALDH1A3+ high
Infection 1
Infection 2
Infection 3
(D)
ALDH1A3+ population (%)
ALDH1A3+ low
ALDH1A3+ med
ALDH1A3+ high
Infection 1
Infection 2
Infection 3
(E)
Taken and added in 1:2 ratio
Administered activating agent
Inhibitor Control Tube
= Activating reagent
= DEAB inhibitor
Incubate
37°C for 45 min
Sample 1A
Sample 1B
Sample 1C
ALDH1+ cells
ALDH1- cells
(F)
ALDH1+ population (%)
Sample 1
Sample 2
Sample 3
(G)
ALDH1+ population (%)
Sample 1
Sample 2
Sample3

Suppl. Fig. 1

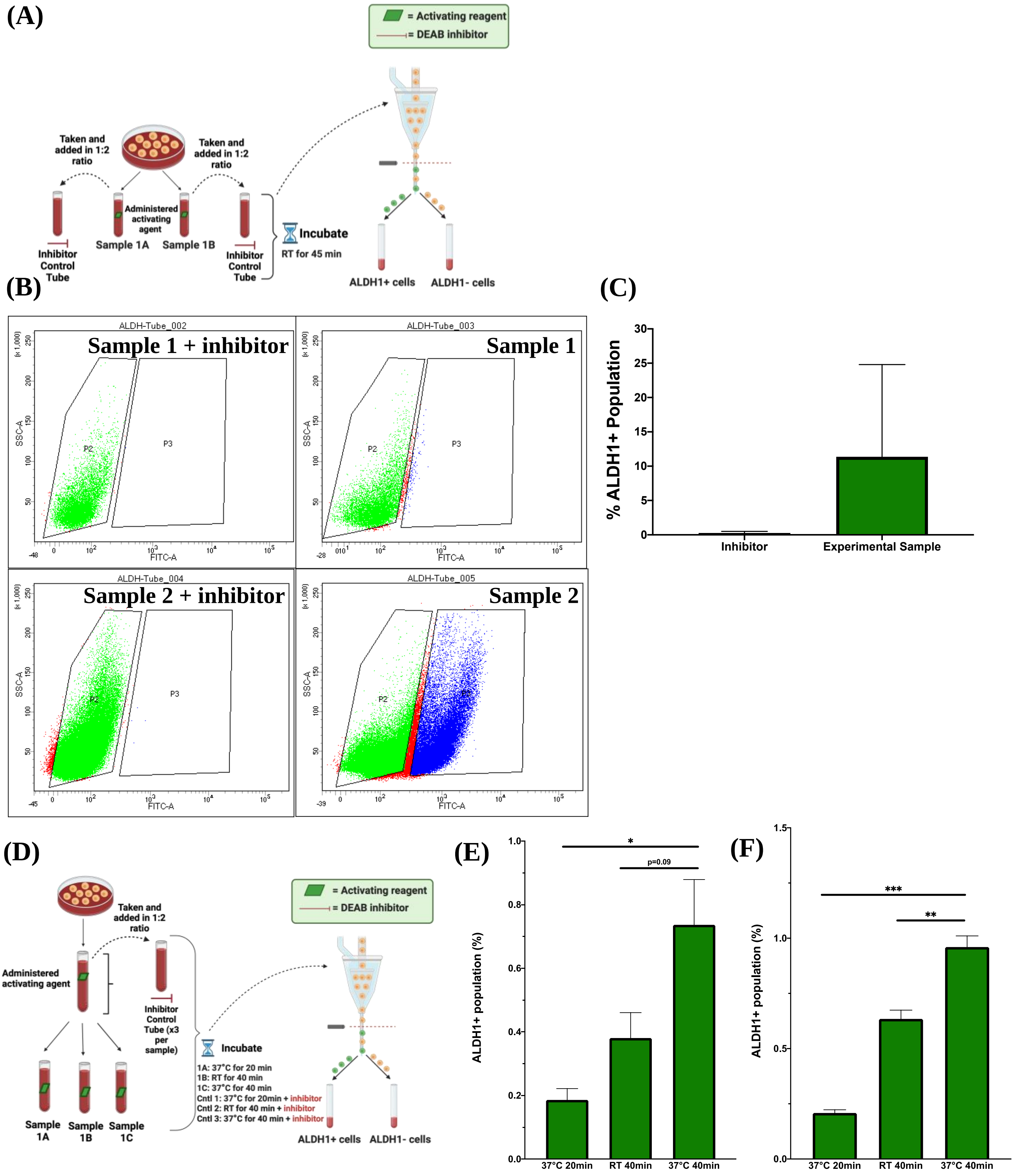
(A)
= Activating reagent
= DEAB inhibitor
Taken and added in 1:2 ratio
Taken and added in 1:2 ratio
Administered activating agent
Sample 1A
Sample 1B
Inhibitor Control Tube
Inhibitor Control Tube
Incubate
RT for 45 min
ALDH1+ cells
ALDH1- cells
(B)
ALDH-Tube_002
Sample 1 + inhibitor
ALDH-Tube_003
Sample 1
ALDH-Tube_004
Sample 2 + inhibitor
ALDH-Tube_005
Sample 2
SSC-A
FITC-A
P2
P3
(C)
% ALDH1+ Population
Inhibitor
Experimental Sample
(D)
Taken and added in 1:2 ratio
Administered activating agent
Inhibitor Control Tube (x3 per sample)
= Activating reagent
= DEAB inhibitor
Incubate
1A: 37°C for 20 min
1B: RT for 40 min
1C: 37°C for 40 min
Cntl 1: 37°C for 20min + inhibitor
Cntl 2: RT for 40 min + inhibitor
Cntl 3: 37°C for 40 min + inhibitor
Sample 1A
Sample 1B
Sample 1C
ALDH1+ cells
ALDH1- cells
(E)
ALDH1+ population (%)
*
p=0.09
37°C 20min
RT 40min
37°C 40min
(F)
ALDH1+ population (%)
***
**
37°C 20min
RT 40min
37°C 40min

Suppl. Fig. 2
